# Butterfly assemblages from Amazonian flooded forests are not more species-poor than from unflooded forests

**DOI:** 10.1101/582742

**Authors:** Rafael M. Rabelo, William E. Magnusson

## Abstract

The Amazonian flooded and upland forests harbour distinct assemblages of most taxonomic groups. These differences can be mainly attributed to flooding, which may affect directly or indirectly the persistence of species. Here, we compare the density, richness and composition of butterfly assemblages in *vaórzea* and *terra firme* forests, and evaluate whether terrain elevation and flooding can be used to predict the assemblage structure. We found that the total abundance and number of species per plot is higher in *vaórzea* than in *terra firme* forests. *Vaórzea* assemblages showed a higher dominance of abundant species than *terra firme* assemblages, in which low-flying Haeterini butterflies had higher abundance. After standardizing species richness by sample size and/or coverage, species richness estimates for *vaórzea* and *terra firme* forests were similar. There was strong turnover in species composition across *vaórzea* and *terra firme* forests associated with terrain elevation, most likely due to differences in the duration of flooding. Despite a smaller total area, less defined vegetation strata, more frequent disturbances and the younger geological age of floodplain forests, Nymphalid butterfly assemblages are not more species poor there than in unflooded forests.

## Introduction

The number and composition of species at a given site is a small subset of the regional species pool because environmental and biotic factors act together or separately to filter species from the regional pool and select the species composition at local scales [1]. Vegetation type is the biotic feature most often used to represent the spatial distribution of forest-dwelling species, and several forests types can be found in Amazonian landscapes.

Upland *terra firme* forests account for approximately 83% of the Amazon basin [2] and are located above the maximum seasonal flood levels of rivers, lakes and large streams. Floodplain *vaórzea* forests, on the other hand, are seasonally flooded by nutrient-rich white- water rivers for 6 to 8 months, and water-level fluctuations can reach up to 14 m [3]. It is estimated that *vaórzea* forests account for ∼ 400,000 km^2^ in the Amazon basin [2].

*Vaórzea* and *terra firme* forests harbour distinct assemblages of trees [4], terrestrial mammals [5], bats [6], birds [7] and litter frogs [8]. These differences in species composition are mainly attributed to flooding, which provides a significant barrier to the persistence of all ground-dwelling and understorey species during the high-water season [9], and even for flying species [6,7]. It has been proposed that *terra firme* has higher species richness than *vaórzea* forest because it offers more niches associated with the understorey vegetation [10]. It is expected that upland forests should contain more speciose assemblages of species groups that can persist in flooded and unflooded forests, since they cover a much larger area [11], have more stratified vegetation [12], suffer less frequent disturbances [13] and have greater geological age [14] than flooded forests. On the other hand, floodplain forests tend to have higher species abundance/biomass [10,15] due to the high forest primary productivity, as the white-water seasonal flooding fertilizes *vaórzea* soils [16].

Butterflies are strongly associated with specific habitats at all life stages [17]. They are relatively sedentary in the larval stage, but are highly vagile in the adult phase and can have seasonal adaptations (phenological or migratory) to environmental changes. Vegetation gradients represent changes in the availability of food resources and physical conditions of the environment, which directly affect the spatial distribution of Amazonian fruit-feeding butterflies [18–20]. Therefore, environmental changes, such as seasonal flooding, may also filter species from the regional pool, affecting local species richness and composition, although no study has been conducted to test that hypothesis.

This study compares the butterfly assemblages of *vaórzea* and *terra firme* forests in a location in Central Amazonia. Specifically, we aim (i) to test whether the density, richness and composition of butterflies differs between *vaórzea* and *terra firme* forests; (ii) to compare the species-abundance distribution between the two forest types; and (iii) to evaluate whether the assemblage-structure pattern is associated with terrain elevation and flooding. We expected to find a higher butterfly density in *vaórzea* forests because they have higher forest primary productivity, which represents higher availability of food resources, than *terra firme*. On the other hand, given that *terra firme* forests represent a more stable environment and cover a larger area, we expected higher species richness in this forest type. Similarly, we predicted that the butterfly assemblage from *vaórzea* forests would have higher dominance of abundant species, and that the species-abundance distribution would be evener in *terra firme* forests. We also expected to find strong turnover in species composition associated with terrain elevation and flooding.

## Materials and Methods

### Study area

Sampling was undertaken near the confluence of Juruaó and Andiraó rivers, in Amazonas State, Northern Brazil (S1 Fig). The interfluvium of the junction of these rivers is protected by the Baixo Juruaó Extractive Reserve [21]. The Juruaó river channel comprises a large floodplain of *vaórzea* forests, which are adjacent to unflooded (*terra firme*) forests. During the high-water season, v*aórzea* forests are flooded by nutrient-rich white-water rivers, with an average annual water-level range of 15 m. Highest river levels occur around May and minima in October [21]. Mean annual temperature and precipitation are around 26 °C and 2255 mm, respectively, with mean precipitation around 60 mm during the dry season [21].

### Sampling design and data collection

Sampling was done in five plots located in *vaórzea* and nine in *terra firme* forests (S1 Fig) at the beginning of the low-water season (July 2018). The sampling design followed the RAPELD method as part of a long-term ecological project that aims to compare the distributions of multiple taxa [22]. Plots (sample units) had 250-m long center lines and were uniformly distributed in the landscape, following the elevation contour to minimize variation in soil conditions and its correlates within the transects [23]. Most plots were separated by at least 1 km from one another, but some *terra firme* plots were separated by only 500 m due to logistical constraints (S1 Fig).

Butterfly surveys were conducted via active and passive sampling. We placed six equally-spaced butterfly baited traps along the center line of each plot. Traps were hung from tree branches in the forest understorey (1.5–2 m high). We baited the traps with a mixture of sugar-cane juice and bananas fermented for 48 h [24] and visited them every 24 h to check for captures and replace the bait. We left the traps active for five consecutive days in each plot. This sampling effort is based on [25], which suggested that it is sufficient to identify ecological responses of understorey fruit-feeding butterfly assemblages.

We also used insect nets to sample low-flying Haeterini species and other Nymphalid species. On each visit to the plots, two researchers with standard 37-cm diameter insect nets actively searched for butterflies during 30 min. All captured individuals were collected for species identification and the specimens were deposited in the Entomological Collection of the Mamirauaó Institute for Sustainable Development, Tefeó, Brazil.

We obtained the elevation data from the digital elevation model (DEM) in the HYDRO1k database developed by the US Geological Survey (http://lta.cr.usgs.gov/HYDRO1K; S1 Fig). We obtained terrain-flooding data from the Synthetic Aperture Radar of the Japonese Earth Resources Satellite – JERS-1 SAR (http://earth.esa.int). JERS-1/SAR images are radar images which, in the Amazon, indicate flooded forests areas by brighter pixels, closed-canopy forests by median brightness, and open water as darker pixels (S1 Fig).

### Data analysis

We compared the total abundance and observed number of species per plot between *vaórzea* and *terra firme* forests with a Kruskal-Wallis test, as the data had a non-normal distribution. We used rarefaction and extrapolation of standardized number of species in order to compare species richness in the two forest types. We standardized the number of species by both number of sampled individuals and sampling coverage, following the recommendations of Chao et al. [26]. Rarefaction and extrapolation were based on sampling coverage in addition to sample size, because standardizing samples by number of individuals usually underestimates species richness of assemblages with more species [27]. We also used the Kolmogorov-Smirnov test to compare the species-abundance curves from the two forest types and sampling methods.

We built a species by site matrix, recording each species (columns) abundance per plot (rows). We standardized the abundances by dividing the number in each matrix cell by the total abundance in the matrix row (plots) to reduce the discrepancy between sites with different numbers of individuals captured. We summarized butterfly species composition by a principal coordinates analysis (PCoA) ordination, based on the Bray-Curtis dissimilarity index. The scores from the first axis derived from this ordination were used to represent the butterfly species composition in each plot. We used a permutational multivariate analysis of variance (PERMANOVA) to evaluate whether the species composition differed between the two forest types. Terrain elevation and flooding were highly correlated (Pearson correlation: *r* = −0.96, *p* < 0.01, S1 Fig). Thus we conducted an analysis of covariance (ANCOVA) to evaluate the effect of elevation on the pattern of assemblage structure, which was represented by first PCoA axis, in each forest type (factor). All analyses were undertaken in the vegan 2.4-4 [28] and iNEXT [29] packages of the R 3.4.4 statistical software [30].

## Results

We captured 357 individuals belonging to 56 butterfly species (S1 Table). The most abundant species in *vaórzea* forests was *Pseudodebis marpessa*, and *Euptychia mollina* was the most abundant in *terra firme*. Singletons and doubletons were represented by 19 species (∼49%) in *vaórzea* forests and 18 (∼67%) in *terra firme*.

The median number of butterflies counted per plot in *vaórzea* forests was 27 (first quartile (Q1) and third quartile (Q3) were 26 and 45, respectively), and was significantly higher than the medium number of butterflies counted in *terra firme* plots (Q1 = 5; median = 9; Q3 = 11; Kruskal-Wallis, *H* = 6.10, *p* < 0.01; Fig 1a). The abundance distribution of species also differed between the two forest types (Kolmogorov-Smirnov, baited traps: *D* = 0.96, *p* < 0.01; insect nets: *D* = 0.67, *p* < 0.01; both methods: *D* = 0.79, *p* < 0.01; Figs 1c and S2). The *vaórzea* assemblage had higher dominance of abundant species (8% of the species made up 50% of all individuals, S3 Fig) than the *terra firme* assemblage, which had an evener distribution of species abundance (19% of the species made up 50% of individuals, S3 Fig).

**Fig 1.**
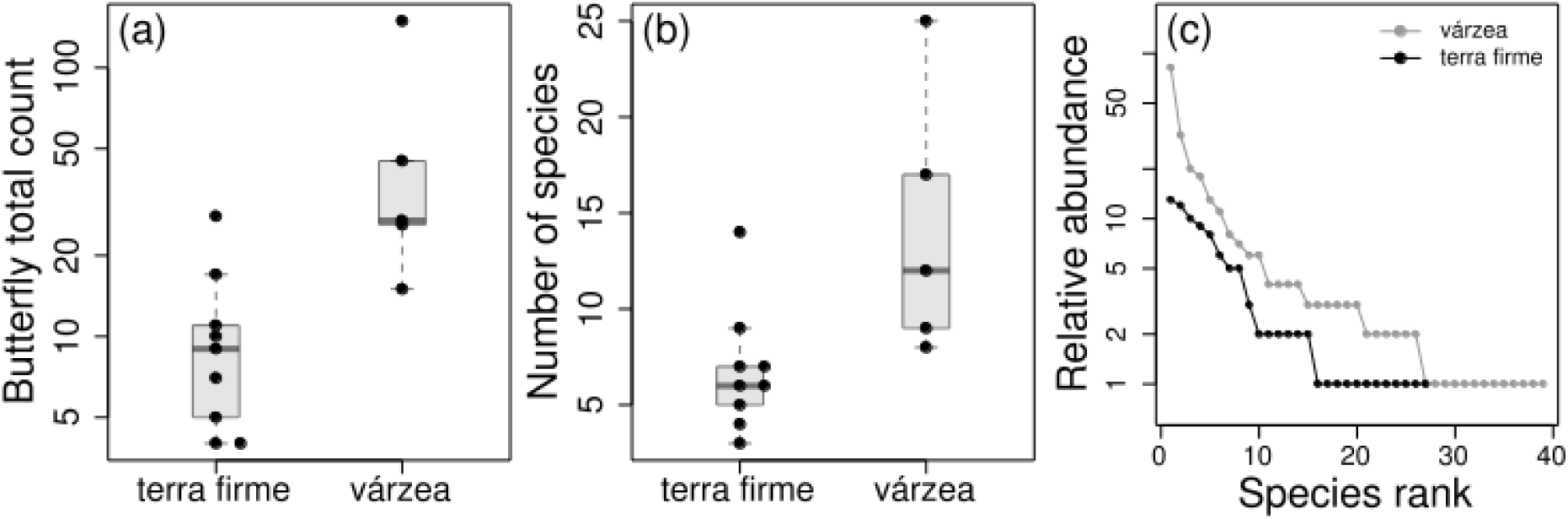
Butterfly counts and number of species in *vaórzea* and *terra firme* forest plots. Difference in butterfly counts (a) and number of species (b) per plot between the two forest types. (c) Assemblage rank-abundance distribution from the two forest types.

The observed number of species per plot was also higher in *vaórzea* than in *terra firme* forests (Kruskal-Wallis, *H* = 5.80, *p* < 0.05; Fig 1b), with a median number of 12 species per plot in flooded forests (Q1 = 9; Q3 = 17) and 6 (Q1 = 5; Q3 = 7) species per plot in upland forests. However, when the species richness estimate was standardized by sample size and coverage, *vaórzea* and *terra firme* forests showed similar species-richness estimates (Fig 2). Although the *terra firme* assemblage had a lower estimated sampling completeness (88%) than *vaórzea* (95%; S4 Fig), the rarefaction and extrapolation of species-richness estimates as a function of sample size or coverage showed similar curves (Fig 2).

**Fig 2.**
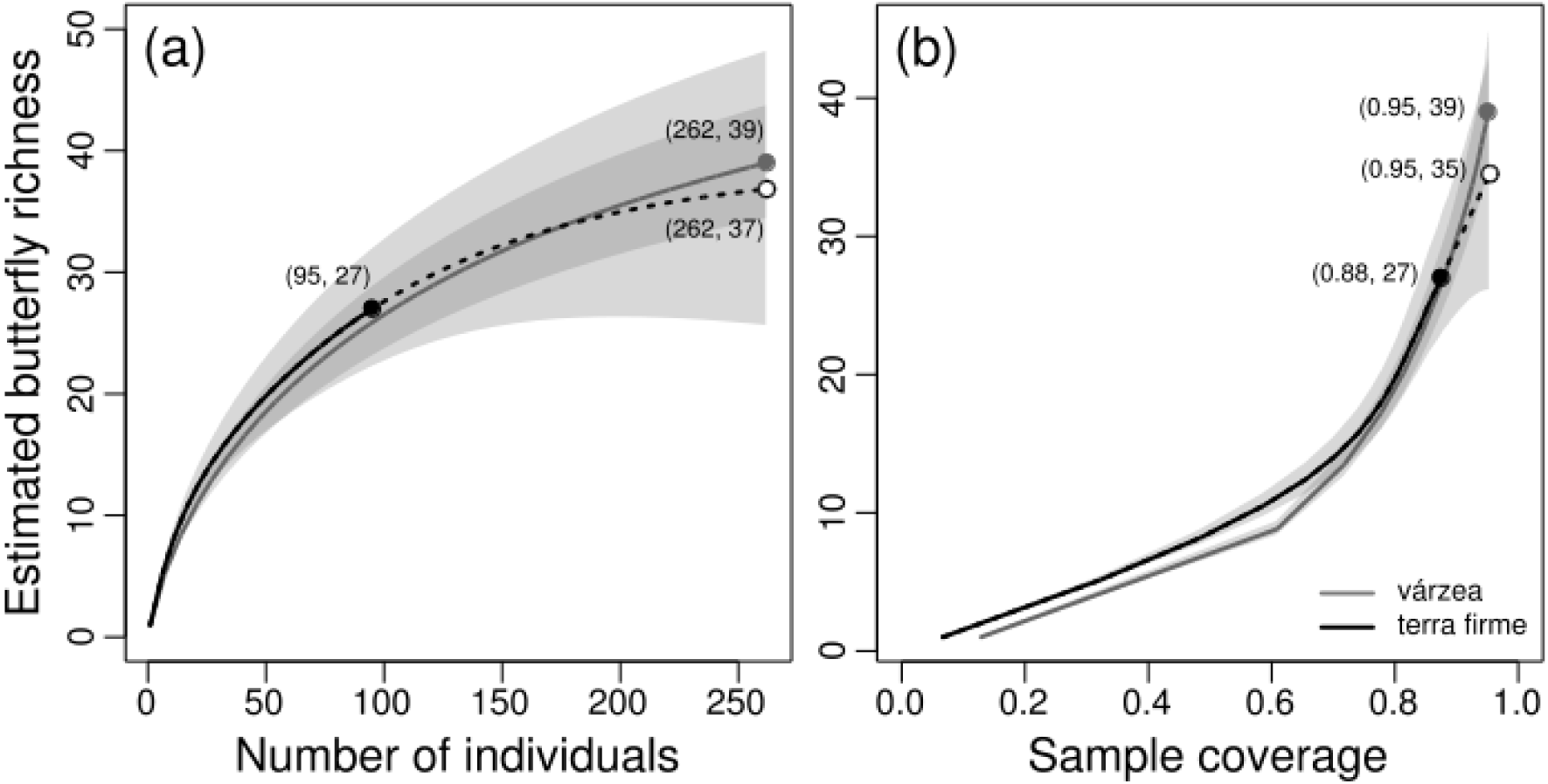
Butterfly richness estimated by rarefaction (solid curves) and extrapolation (dashed curves) based on sample size (a) and completeness (b), with corresponding 95% confidence intervals (shaded areas). Solid circles indicate the observed species richness and open circles indicate the extrapolated richness in *terra firme* assemblages based on number of individuals (a) or sample coverage (b). Numbers within parentheses indicate the coordinates of points in both graphs. Although estimated richness in *vaórzea* seemed to be slightly higher than *terra firme* at its maximum sample size (262 individuals in “a”) or completeness (0.95 of coverage in “b”), the confidence intervals overlap and indicate the there is no statistically significant difference in richness between the two forest types.

The PCoA ordination of plots along the two first axes explained 42% of the variation in species composition. There was a marked difference between butterfly composition of *vaórzea* and *terra firme* forests (PERMANOVA, F = 4.23, *p* < 0.01), captured mainly by the first axis (Fig 3a) due to the strong turnover of species composition between forest types (Fig 3b). The *vaórzea* species composition was not a nested subset of the *terra firme* assemblage. The change in species composition was associated with forest types (F = 19.22; *p* < 0.01), but without effect of terrain elevation within each forest type (F = 1.27; *p* = 0.29; Fig 3c).

**Fig 3.**
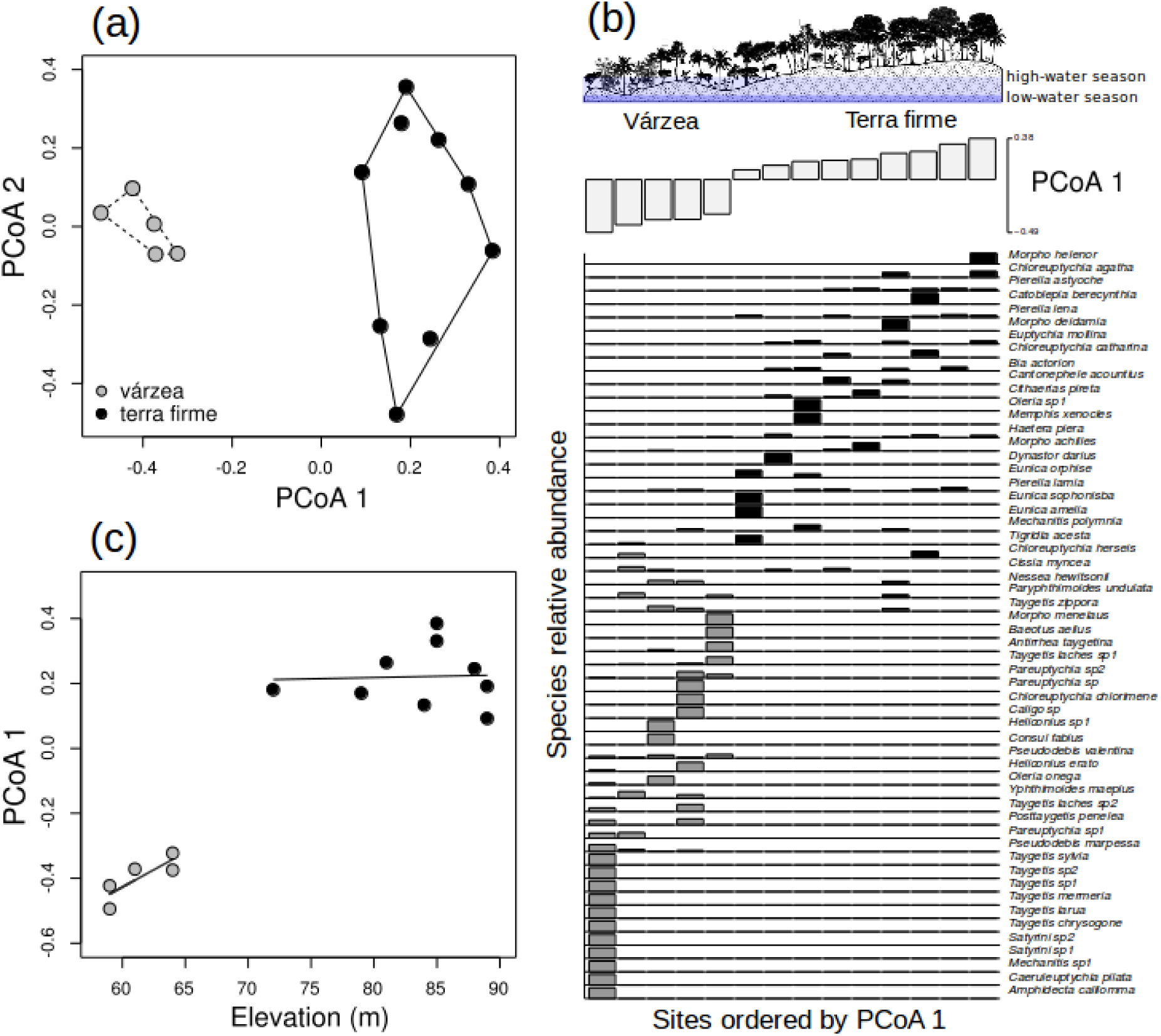
Changes in species composition between *vaórzea* and *terra firme* forests. (a) Similarity in butterfly species composition of plots represented by the distances formed in the two axes derived from the PCoA ordination. Each point in the graph represent a plot located in *vaórzea* or *terra firme* forest and the distance between points represents the similarity of plot in terms of species composition. (b) Distribution of butterflies across sample sites. Sample sites are ordered by the first PCoA axis and bar heights show the relative abundance of butterfly species across *vaórzea* (gray) and *terra firme* (black) plots. (c) Change in species composition (PCoA 1) with elevation within each forest type.

## Discussion

We found higher butterfly total density in *vaórzea* than in *terra firme* forests, which is the same pattern reported in studies of bats [10] and primates [15]. The higher density of herbivorous, frugivorous and nectarivorous species (such as butterflies, primates and frugivorous bats) in *vaórzea* forests is probably due to the higher availability of food resources for these species. Seasonal flooding by white-water rivers provides an extra input of nutrients in *vaórzea* soils, which increases forest primary productivity [16]. Bobrowiec et al. [6] found that the abundance of frugivorous bats in *vaórzea* forests is even higher during the high-water season. However, for Amazonian fruit-feeding butterflies, adults tend to be more abundant during the early and mid dry season, and less abundant during the wet season [31], when they probably occur in other life stages, such as herbivorous caterpillars.

We found that *vaórzea* forests had a higher number of species per plot (i.e., higher species density) than *terra firme*. This apparent difference in the number of butterfly species between the two forest types occurs because we sampled a much higher number of individuals per plot in *vaórzea* forest. Therefore, the difference in the amount of nutrients between the two forest types [16] may also explain the difference in the species density between *vaórzea* and *terra firme* forests. However, the higher number of species per plot found in *vaórzea* forests did not result in a higher total butterfly richness in the flooded forest.

The *terra firme* assemblages had a lower sampling completeness than *vaórzea* forest, despite the larger survey effort (nine surveyed plots), and a higher proportion of rare species (singletons and doubletons). When extrapolating the *terra firme* species richness to the same size/coverage as the *vaórzea*’s sample, we found that both assemblages showed similar rarefaction and extrapolation curves (Fig 2), indicating that they have similar overall richness.

Poorer assemblages in *vaórzea* have been consistently documented for several animal groups [5,6,15], and seasonal inundation is the potential explanation for the lower number of terrestrial and understorey species. However, few studies have attempted to estimate species richness by standardizing the number of species by sample size/coverage prior to undertaking such comparisons (but see [10]). A comparison of bat assemblages between these two forest types, found a higher bat richness in *terra firme* than in *vaórzea*, and the authors suggested that the higher richness occurs because upland forests contain more niches associated with the understorey vegetation [10]. The higher complexity in the *terra firme* forest structure [16] may also increase the diversity of niches to be occupied by butterflies, explaining the similarity in species richness between the two forest types, despite the lower abundance in the upland forest.

Three butterfly species made up 50% of all individuals from the *vaórzea* assemblages: *Pseudodebis marpessa, Oleria onega* and *P. valentina* (S3 Fig). Oviposition of *Pseudodebis* species generally occurs in May–June and its life cycle lasts around 50 days [32], which may explain the high abundance we found during our survey (July). Additionally, *Pseudodebis* species feed on the bamboo *Guadua angustifolia* [32], locally known as *“taboca”*, which was highly abundant the *vaórzea* plot where we surveyed most *Pseudodebis* butterflies (R. Rabelo, person. obs.). *Oleria* are Ithomiinae butteflies that are known to feed on alkaloid-rich host plants, which make the adults unpalatable to predators and all species are engaged in mimicry [33,34]. Although adults are unpalatable, it has been suggested that their eggs may be subject to predation or removed from leaves by *Ectatomma* ants, which are often found on *Solanum* species [35]. As *Ectatomma* ants are weak swimmers [36] and do not normally occur in Amazonian seasonally-flooded forests [37], we hypothesize that their absence may favor the high abundance of *Oleria* in *vaórzea* forests.

The rank-abundance distribution was slightly evener in the *terra firme* assemblage, with five species (19%) summing more than 50% of all individuals from the upland assemblage (S3 Fig). *Euptychia molina* was the most abundant species in *terra firme* assemblage, followed by three species from the Haeterini tribe. *Euptychia* butterflies are known for their strong relationship with their host plants, which are among the oldest plant lineages: Selaginellaceae (Lycopsidophyta) and Neckeraceae (Bryophyta) [38,39]. These plant lineages are often obligate terrestrial (*Selaginella*) and do not occur in floodplain forests [40,41], which may be the reason why *E. molina* was abundant and restricted to *terra firme*.

The evener rank-abundance distribution in *terra firme* forests was mainly caused by the Haeterini butterflies, which tended to be more abundant in this forest type (S1 Table). Three of five Haeterini species were restricted to this forest type (*Cithaerias pireta, Pierella astyoche* and *P. lena*, Fig 2b). Haeterini butterflies are low-flying ground-dwelling species that feed mainly on rotting fruits and other decaying material on the forest floor [42], and adults can be abundant throughout the year [43]. The host plants for these species are *Spathiphyllum* sp. for *Haetera* butterflies [44], *Philodendron* sp. for *Cithaerias* butterflies [45] and mainly species from Heliconiaceae and Maranthaceae for *Pierella* butterflies [38]. *Spathiphyllum* and *Phylodendron* species do not occur in *vaórzea* forests, and terrestrial species of Heliconiaceae and Maranthaceae may occur in inundated forests, although they are not usually common [40]. Therefore, the seasonal flooding of *vaórzea* forests may explain the higher abundance and constrained distribution of Haterini butterflies and their host plants to *terra firme* forests.

We found a pronounced difference in butterfly species composition between *vaórzea* and *terra firme* forests. The strong turnover of species across forest types was captured by the first PCoA axis. We have discussed some examples of how *vaórzea* flooding can affect butterflies and their host plants distribution through increased soil fertility and, consequently, forest primary productivity [16], which results in differences in resources availability – soil nutrients that are resources for host plants, which in turn are resources for butterflies. Also biotic constraints due to interaction with predators (e.g., *Ectatomma* ants that prey upon *Oleria* eggs and their host plants [35]); and flooding *per se*, which constrains the distribution of low-flying Haeterini butterflies (and several host plant species) to *terra firme* forests. Therefore, the results of this study suggest that environmental and biotic filters override the effects of vegetation stratification and effects of source area on differences in the composition of butterfly assemblages in flooded and unflooded Amazonian sites at local scales.

## Acknowledgements

This research was funded by the Instituto de Desenvolvimento Sustentaóvel Mamirauó (IDSM-OS/MCTI) and by the Gordon and Betty Moore Foundation. We thank the logistical support from Instituto de Desenvolvimento Sustentaóvel Mamirauaó and from Program for Biodiversity Research (PPBio) and the National Institute for Amazonian Biodiversity (INCT-CENBAM; FAPEAM/FDB/INPA, #003/2012; and CNPq, #573721/2008-4, #722069/2009). Butterfly specimens were collected under permisson SISBIO 57444. We thank Iury Valente e Geanne Pereira from Departamento de Mudanças Climaóticas e Unidades de Conservaçaão (DEMUC) for providing butterfly sampling equipment and Joaão Valsecchi for insightful contributions in the early ideas of this work. We also thank the Cumaru village for their cordial reception.

## Supporting Information

**S1 Fig.** Distribution of sample plots in *vaórzea* and *terra firme* forests. (a) Terrain elevation and (b) flooded areas. (c) Correlation between elevation and flooding at sample plot locations.

**S1 Table**. Abundance of Nymphalidae butterflies collected in 14 plots (five in *vaórzea* and nine in *terra firme* forests) in Baixo-Juruaó Extractive Reserve, Amazonas State, Brazil.

**S2 Fig.** Species-abundance distribution of butterfly species in vaórzea and terra firme forests sampled with baited traps (left) and insect nets (right). In both sampling methods, the rank-abundance curves of species for different habitats were found to come from different distributions (Kolmogorov-Smirnov, baited traps: *D* = 0.96, *p* < 0.01; insect nets: *D* = 0.67, *p* < 0.01).

**S3 Fig.** Rank-abundance distribution of butterfly species in *vaórzea* and *terra firme* forests.

**S4 Fig.** Plot of sample coverage for rarefied samples (solid line) and extrapolated samples (dashed line) as a function of sample size for butterfly samples from *vaórzea* and *terra firme* forests, with 95% confidence intervals (shaded areas). Observed samples are denoted by filled circles. Each of the two curves was extrapolated up to double its observed sample size. The numbers in parentheses are the sample size and the estimated sample coverage for each reference sample. Unfilled circles represent the number of individuals to be sampled from each assemblage when sample coverage is 0.954 (i.e., the sample coverage at double the observed sample size for the *terra firme* assemblages).

